# Influence of GST- and P450-based metabolic resistance to pyrethroids on blood feeding in the major African malaria vector *Anopheles funestus*

**DOI:** 10.1101/2020.03.16.993535

**Authors:** Lynda Nouage, Emmanuel Elanga-Ndille, Achille Binyang, Magellan Tchouakui, Tatiane Atsatse, Cyrille Ndo, Sévilor Kekeunou, Charles S. Wondji

**Author notes:** These authors contributed equally to this work (Lynda Nouage, Emmanuel Elanga-Ndille). Corresponding authors: Emmanuel ELANGA-NDILLE (;) and Lynda NOUAGE.

## Abstract

Insecticide resistance genes are often associated with pleiotropic effects on various mosquito life-history traits. However, very little information is available on the impact of insecticide resistance, especially metabolic resistance, on blood feeding process in mosquitoes. Here, using two recently detected DNA-based metabolic markers in the major malaria vector, *An. funestus*, we investigated how metabolic resistance genes could affect blood meal intake.

After allowing both field F1 and lab F8 *Anopheles funestus* strains to feed on human arm for 30 minutes, we assessed the association between key parameters of blood meal process including, probing time, feeding duration, blood feeding success and blood meal size, and markers of glutathione S-transferase (*L119F-GSTe2*) and cytochrome P450 (*CYP6P9a*_R) - mediated metabolic resistance. None of the parameters of blood meal process was associated with *L119F-GSTe2* genotypes. In contrast, for *CYP6P9a*_R, homozygote resistant mosquitoes were significantly more able to blood-feed than homozygote susceptible (OR = 3.3; CI 95%: 1.4-7.7; P =0.01) mosquitoes. Moreover, the volume of blood meal ingested by CYP6P9a-SS mosquitoes was lower than that of CYP6P9a-RS (P<0.004) and of CYP6P9a-RR (P<0.006). This suggests that *CYP6P9a* gene affects the feeding success and blood meal size of *An. funestus*. However, no correlation was found in the expression of *CYP6P9a* and that of genes encoding for salivary proteins involved in blood meal process.

This study suggests that P450-based metabolic resistance may increase the blood feeding ability of malaria vectors and potential impacting their vectorial capacity.

## Introduction

Despite significant progress made since 2000s in reducing its burden in sub-Sahara Africa, malaria remains a major public health burden in this region [1]. This disease is caused by a *Plasmodium* parasite transmitted by *Anopheles* mosquito species while taking blood meal on humans. Blood feeding, is essential for female mosquito’s fecundity [2], as *Anopheles* species like all anautogenous female mosquitoes, require blood meal to obtain amino acids needed to synthesize yolk proteins for eggs maturation [3,4].

Mosquito’s blood feeding success is facilitated by the pharmacological proprieties of salivary gland proteins [5]. Indeed, some salivary proteins such as anopheline antiplatelet protein (AAPP), apyrase, gSG6 and members of D7 family have been identified as vasodilators, anticoagulants and inhibitors of platelet aggregation allowing mosquitoes to overcome host haemostatic mechanisms and to have a successful blood meal [5–8].

Mosquito’s fecundity was shown to vary with source and size of the blood meal with difference of these two parameters resulting in significant variations of the number of eggs laid by each female mosquito [4,8]. It has been shown that the number of eggs laid per female is positively associated to the amount of blood ingested as, larger blood meals resulted in the increase of the number of female that developed eggs and the number of eggs per female [9,10]. The volume of blood meal of a mosquito could be affected by a range of intrinsic (host immunity) and extrinsic factors including ambient temperatures, mosquito age, parity status, gonotrophic cycle, blood feeding history and infection status [10]. More recently, it was reported that exposure to pyrethroids could also significantly influence the blood meal process and the blood meal volume ingested by *Anopheles gambiae* mosquito [11].

Pyrethroids (PY) are the insecticide class mostly used in the last two decades through ITNs and IRS strategies to control malaria transmission [12]. Unfortunately, the widespread use of these insecticides has favoured the development of resistance in malaria vector species [13,14]. Resistance to pyrethroids involves two main mechanisms: (i) metabolic resistance, due to the increase expression level of detoxifying enzymes, belonging to three families: the cytochrome P450 monooxygenases, the glutathione S-transferases and the carboxyesterases; and (ii) target-site resistance due to mutations in the voltage sodium channels known as knock-down *(kdr)* mutations [15,16]. Although resistance mechanisms help mosquitoes survive under continuous insecticide pressure, these actions are costly and may negatively affect mosquito’s fitness including body size, adult longevity, larval development time, fecundity, fertility, mating competitiveness and blood feeding capability [17–19]. For target-site resistance, a decreased longevity and increased larval development time have been reported in *kdr*-pyrethroid-resistant mosquitoes [20,21]. Moreover, a recent study suggested that *kdr*-based resistance could impact blood feeding with heterozygote (*kdr*-RS) and susceptible (*kdr*-SS) mosquitoes taking higher blood volume than homozygote (*kdr*-RR) resistant individuals [11]. In contrast, little is known on the impact of metabolic resistance as DNA-based markers were not previously available for this mechanism preventing to investigate its physiological impact on the blood feeding process in mosquitoes. However, taking advantage to the identification of the first DNA-based metabolic marker in *An. funestus* mosquito, one study reported that a GST-based metabolic resistance caused by a leucine to phenylalanine amino acid change at codon 119 in the glutathione S-transferase epsilon 2 *(L119F-GSTe2)* [22], has a detrimental impact on *An. funestus* fitness as field mosquitoes exhibited a reduced fecundity and slower larval development but an increased adult longevity [23]. On the other hand, a new DNA-based assay was recently designed for cytochrome P450-mediated resistance (the *CYP6P9a-R* in *An. funestus*. This marker used to show that mosquitoes carrying this P450-resistant allele survived and succeeded in blood feeding more often than did susceptible mosquitoes when exposed to insecticide-treated nets [24]. The design of assays for both GST- and P45O-based resistance now offers a great opportunity to explore how the blood feeding process is impacted by metabolic resistance mechanism in malaria vectors and further assess how resistance may impact the vectorial capacity of mosquitoes to transmit malaria in the field.

Here, we assessed the effect of metabolic resistance to pyrethroids on the blood feeding process in *An. funestus*, using the two DNA-based metabolic resistance markers: *L119F-GSTe2* and *CYP6P9a*-R [22,24]. Specifically, we assessed the association between the genotypes of these metabolic resistance markers and key parameters of blood feeding including mosquito probing time, feeding duration and the bloodmeal size revealing a contrasting impact.

## Material and Methods

### Mosquito collection and rearing

Experiments were carried out using both field and lab strains of *An. funestus*. Field mosquitoes (F_1_) were generated from indoor resting female (F0) collected in Mibellon (6°46’N, 11^0^ 70’E), a village located in rural area in the savanna-forest region in Cameroon, Central Africa where the *L119F-GSTe2* has previously been detected [25]. Blood-fed field collected females were kept in paper cups and transported to the insectary of the Centre for Research in Infectious Diseases (CRID) in Yaoundé where they were kept for 4-5 days until they became fully gravid and then induced to lay eggs using the forced eggs-laying method [26]. The eggs were put in paper cups containing water to hatch, after which the larvae were placed in trays and reared to adults. To assess the effect of *CYP6P9a* marker absent in Cameroon [24], F_8_ progenies were generated from reciprocal crosses established between the pyrethroid susceptible laboratory strain (FANG) and the resistant (FUMOZ-R) lab strain. These two *An. funestus* lab strains were colonized from mosquitoes collected in Southern Africa region. FUMOZ is a pyrethroid resistant strain established in the insectary in 2000 from wild-caught *An. funestus* mosquito species from southern Mozambique [27]. Previous study reported that the over-expression of two duplicated P450 gene, *CYP6P9a* and *CYP6P9b*, is the main mechanism driving pyrethroid resistance in this strain [28,29] for which the *119F-GSTe2* allele is absent [22]. The FANG strain is a completely susceptible to pyrethroids colonized from Calueque in southern Angola in 2002 [27].

### Blood feeding experiments and blood meal size quantification

#### Blood feeding process

As blood meal volume has previously been reported to correlate with mosquito size [2], individuals used for blood feeding experiments were first grouped according to their size. Mosquito size was determined by weighing (using an analytical micro-scale, SARTORIUS, Goettingen, Germany) each individual (adult females aged 3-7 days) starved for 24h and immobilized by chilling them for 2 minutes at 5°C. Each mosquito was then transferred in paper cups and was kept in the cups for about an hour before been given a blood meal. Mosquitoes were allowed to bite for 30 min on the bare forearm of a single human volunteer after an informed consent.

The duration of probing and blood feeding was assessed using a batch of 120 F_1_ female field-collected mosquitoes. For this purpose, mosquitoes were individually transferred in polystyrene plastic cups covered with net. They were allowed to rest for 15 min before observations began. During the blood intake, each mosquito was filmed using a Digital HD Video Camera (Canon PC2154, Canon INC, Japan) placed beside the plastic cup. At the end of the time allowed for feeding, the film for each mosquito was analysed and the parameters such as probing time (defined as the time taken from initial insertion of the mouthparts in the skin until the initial engorgement of blood [5] and total feeding duration, were recorded, using a digital timer. For the lab strain mosquitoes, because of low density of female mosquitoes obtained at F_8_ generation from reciprocal crosses, we were not able to run experiments to estimate the probing and the feeding duration of this strain.

To determine the blood meal size, for both strains, batches of 25 mosquitoes grouped according to their weight were allowed to bite on human arm. In this case, neither the probing time nor the feeding duration were recorded. After the trial, whole abdomen of successfully fed mosquitoes (evidenced by red-coloration engorgement of the abdomen) was extracted and stored in an individual 1.5 ml microtube at – 20°C to measure the blood meal size. The rest of the carcasses as well as unfed mosquitoes were kept individually in microtube containing RNA-later and stored at −20°C.

### Blood meal size quantification

The volume of blood ingested by each mosquito was determined by quantifying the haemoglobin amount, as previously described [30]. Briefly, abdomen of blood fed mosquitoes were homogenized in 0.5 ml of Drabkin’s reagent which converts the haemoglobin into haemoglobin cyanide (HiCN). After 20 minutes at room temperature and the addition of 0.5 ml of chloroform solution, samples were centrifuged at 5600 rpm for 5 min. The aqueous supernatant containing HiCN was transferred in a new 1.5 ml microtube. An aliquot of 200μl from each sample was transferred to a microplate and the optical density (OD) read at a wavelength of 620nm in a spectrophotometer (EZ Read 400, biochrom, Cambridge, UK). OD for each sample were read in duplicate and the average value between the two replicates was considered as OD value of the sample. In parallel, OD read on various amounts of human volunteer blood added to Drabkin’s reagent in individual microtubes were used as control to generate calibration curves and the regression line used to assess the relationship between OD and blood volume. For each sample, the blood meal size was estimated according to the weight by dividing the blood volume estimated using the regression line by the average weigh of each batch of mosquitoes constituted after the weighing. The blood meal size was then expressed in μL of blood per mg of weight.

### Molecular species identification

Genomic DNA (gDNA) was extracted from both blood-fed and unfed mosquitoes using the Livak protocol [31]. Instead of using the whole body as done for unfed mosquitoes, DNA was extracted from the carcasses of fed mosquitoes after removing the abdomen for blood volume quantification. The concentration and purity of the extracted gDNA were subsequently determined using NanoDrop^™^ spectrophotometer (Thermo Scientific, Wilmington, USA) before storage at −20 °C. An aliquot of gDNA extracted from field-collected strain was used for molecular identification through a polymerase chain reaction [32] to determine species composition of *An. funestus* group among our samples.

### Genotyping of *L119F-GSTe2* mutation in field-collected strain

The *L119F-GSTe2* mutation was genotyping using gDNA extracted from carcasses of field-collected strain and following allele specific PCR diagnostic assay previously described [23]. The primers sequences are given in table S1. PCR was performed in Gene Touch thermalcycler (Model TC-E-48DA, Hangzhou, 310053, China), in a reaction volume of 15 μl using 10 μM of each primer, 10X Kapa Taq buffer A, 0.2 mM dNTPs, 1.5 mM MgCl2, 1U Kapa Taq (Kapa Biosystems, Wilmington, MA, USA) and 1μl of genomic DNA as template. The cycle parameters were: 1 cycle at 95 °C for 2 min; 30 cycles of 94 °C for 30 s, 58 °C for 30 s, 72 °C for 1 min and then a final extension at 72 °C for 10 min. PCR products were separated on 2% agarose gel by electrophoresis. The size of the diagnostic band was 523 bp for homozygous resistant (RR) and 312 bp for homozygous susceptible (RS), while heterozygous (SS) showed the two bands.

### Genotyping of *CYP6P9a-R* allele in lab strain mosquitoes

The *CYP6P9a* resistance marker was genotyped using the protocol recently designed by [24]. PCR-RFLP were carried out using gDNA extracted from the carcasses of individual of F_8_ generation from reciprocal crosses between FANG and FUMOZ strains used for blood feeding. Briefly, a partial *CYP6P9a* upstream region was amplified in a final volume of 15μl PCR mixture contained 1.5μl of 10X Kapa Taq buffer A (Kapa Biosystems, Wilmington, MA, USA), 0.12μl of 5 U/μl KAPA taq, 0.12μl of 25μM dNTP, 0.75μl of 25μM MgCl2, 0.51μl of each primer, 10.49μl of dH2O and 1μl of genomic DNA. The PCR cycle parameters were as followed: the initial denaturation step at 95°C for 5 minutes followed by 35 cycles of 94°C for 30 seconds, 58°C for 30 seconds and 72°C for 45 seconds and a final extension step of 72°C for 10 minutes. The PCR products were size separated on a 1.5 % agarose gel stained with Midori Green Advance DNA Stain (Nippon genetics Europe GmbH) and visualised using a gel imaging system to confirm the product size (450bp). Then, the PCR product was incubated at 65°C for 2 hours. This was done in 0.2ml PCR strip tubes using 5μl of PCR product, 1μl of cutSmart buffer, 0.2μl of 2 units of Taq1 enzyme (New England Biolabs) and 3.8μl of dH20. Size separation was done on a 2.0% agarose gel stained with Midori Green Advance DNA Stain at 100 V for 30 minutes. The gel was visualised using the gel imaging system.

### Gene expression profiling of major salivary genes encoding proteins involved in blood meal process

The expression profiles of a set of salivary genes encoding for proteins involved in blood meal process was compared between CYP6P9a-RR, CYP6P9a-RS and CYP6P9a-SS *An. funestus* mosquitoes. For each gene, two pairs of exon-spanning primers was designed for each gene using Primer3 online software (v4.0.0; http://bioinfo.ut.ee/primer3/) and only primers with PCR efficiency between 90 and 110% determined using a cDNA dilution series obtained from a single sample, were used for qPCR analysis. Taking into account this criteria of efficiency, only the AAPP and four members of D7 family genes (D7r1, D7r2, D7r3, and D7r4) were used for this analysis. Primers are listed in Table S1. Total RNA was extracted from three batches of 10 whole females of 3-5 days old from CYP6P9a-RR, CYP6P9a-RS and CYP6P9a-SS mosquitoes. RNA was isolated using the RNAeasy Mini kit (Qiagen) according to the manufacturer’s instructions. The RNA quantity was assessed using a NanoDrop ND1000 spectrophotometer (Thermo Fisher) and 1μg from each of the three biological replicates for each batch of mosquitoes was used as a template for cDNA synthesis using the SuperScript III (Invitrogen, Waltham, Massachusetts, USA) with oligo-dT20 and RNase H, following the manufacturer’s instructions. The qPCR assays were carried out in a MX 3005 real-time PCR system (Agilent, Santa Clara, CA 95051, United States) using Brilliant III Ultra-Fast SYBR Green qPCR Master Mix (Agilent). A total of 10 ng of cDNA from each sample was used as template in a three-step program involving a denaturation at 95 °C for 3 min followed by 40 cycles of 10 s at 95 °C and 10 s at 60 °C and a last step of 1 min at 95 °C, 30 s at 55 °C, and 30 s at 95 °C. The relative expression and fold-change of each target gene in CYP6P9a-RR and CYP6P9a-RS relative to CYP6P9a-SS was calculated according to the 2^-ΔΔCT^ method incorporating PCR efficiency after normalization with the housekeeping RSP7 ribosomal protein S7 (VectorBase ID: AFUN007153;) and the actin 5C (vectorBase ID: AFUN006819) genes for *An. funestus*.

### Statistical analysis

All analyses were conducted using GraphPad Prism version 7.00 software. We estimated a Fisher’s exact probability test and the odds-ratio of *L119F-GSTe2* and *CYP6P9a* genotypes (homozygous resistant = RR, heterozygote resistant=RS and homozygous resistant =SS) and both susceptible (S) and resistant (R) alleles. This allowed to assess the association between: a) insecticide resistance and mosquito’s weight by comparing the proportions of the genotypes of both genes in each group established after weighing; b) blood feeding success and insecticide resistance by comparing the proportion of each genotype in both fed and unfed mosquitoes. After arbitrary regrouping the time into four different classes with same amplitude, the duration of probing and feeding was analysed with chi-square test by comparing the proportion of *L119F-GSTe2* and *CYP6P9a* genotypes in each defined time class. After estimating the median of weighted blood meal for each genotype, Kruskal-Wallis and Mann-Whitney tests were used to compare difference between more than two groups and between two groups, respectively.

## Results

### Metabolic resistance genes and *An. funestus* mosquito’s weight

A total of 1,200 and 273 female mosquitoes were weighted, respectively for field strain (F1 generation) and lab strain (F_8_ generation). The mean weight of a mosquito was 0.9 ± 0.010 mg (minimum = 0.2 mg; maximum = 2.3mg) and 0.89 ± 0.016 mg (minimum = 0.2 mg; maximum = 1.7mg) for field and lab strain respectively. No significant difference was found between the mean weights of two strains. For all the analyses, we arbitrarily grouped mosquitoes according to their weight values, into two different classes as followed: [0 - 1.0] mg and [1.1 - 2.4] mg. Analysis of the distribution of *L119F-GSTe2* mutation genotypes in each class of field strain mosquitoes showed no association between the mosquito’s weight and *L119F-GSTe2* genotypes (χ^2^ = 0.15;*p* = 0.9; OR =1.2, 95%CI: 0.3742 - 4.176, for RR *vs* RS; OR=1.1, 95%CI: 0.3659 - 3.606 for RR *vs* SS; OR = 0.9, 95%CI: 0.4943 - 1.709 for RS *vs* SS) (Figure 1 and Table 1). This absence of correlation between the *L119F-GSTe2* genotypes and the weight of mosquitoes was confirmed at the allele level (OR=1; 95%: CI: 0.5–2.0; *p* =0.5) showing that the L119F mutation may not impact the weight of this *An. funestus* field population (Table 1). In contrast, a significant association was observed between *CYP6P9a* genotypes and the weight of mosquito (χ^2^= 29.54, p<0. 0001). Indeed, proportions of RR and RS genotypes were higher than that of SS in the lowest weight class, whereas, for larger weight, mosquitoes with SS genotype were more abundant (67.2%). This association is further supported by odds ratio estimates showing that proportions of homozygote resistant (RR) (OR=5.4; CI 95%: 2.3-12.7; *p*<0.0001) and heterozygote (RS) (OR=5.6; CI 95%: 2.8-11.1; *p*<0.0001) mosquitoes are significantly higher in lowest weight class than the larger one when compared to homozygote susceptible mosquitoes (Table 1). Overall, mosquitoes harbouring the *CYP6P9a*-S susceptible allele displayed higher weight compared to those with the *CYP6P9a*-R resistant allele (OR=2.8; CI 95%: 1.5–5.0; p =0.0003 (Table 1) suggesting that over-expression of the *CYP6P9a* gene is reducing the weight of pyrethroid resistant *An. funestus* mosquitoes.

**Figure 1:**
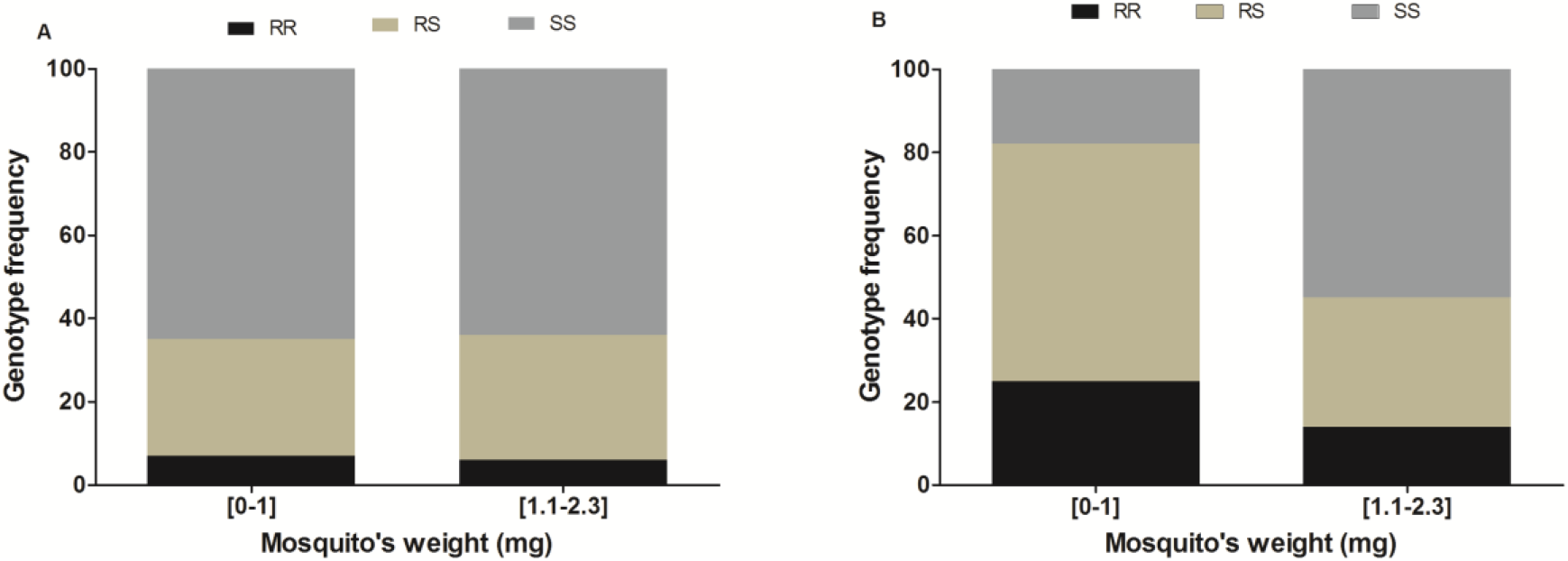
Distribution of genotypes of *L119F-GSTe2* (A) and *CYP6P9a*-R (B) markers according to *An. funestus* mosquito weight.

**Table 1:** level of association of *L119F-GSTe2* and *CYP6P9a*-R genotypes with mosquito weight by comparing low (0-1.0mg) and high (1-2.4mg) weight samples.

| Genotypes | *L119F-GSTe2* |  | *CYP6P9a*-R |  |
| --- | --- | --- | --- | --- |
| | Odds ratio | $p$-value | Odds ratio | $p$-value |
| RR vs SS | 1.1 (0.4-3.6) | 0.5 | 5.4 (2.3-12.7) | < 0.0001 |
| RS vs SS | 0.9 (0.5-1.7) | 0.4 | 5.6 (2.8-11.1) | < 0.0001 |
| RR vs RS | 1.2 (0.4-4.1) | 0.5 | 1.0 (0.5-2.3) | 0.5 |
| S vs R | 1 (0.5-2.0) | 0.5 | 2.8 (1.5-5.0) | 0.0003 |

SS: homozygote susceptible; RR: homozygote resistant; RS: heterozygote; \* significant difference $p < 0.05$ .

### Impact of *L119F-GSTe2* and *CYP6P9a* mutations on *An. funestus* blood feeding success

#### L119F-GSTe2

out of the 1,200 individuals from field strain mosquitoes that were allowed to take a blood meal, 457 (39.6%) successfully fed whereas 743 did not. Among blood-fed mosquitoes, a total of 360 were successfully genotyped and 7% (24/360) were homozygote resistant (RR), 28% (103/360) were heterozygotes resistant (RS) and 65% (233/360) were homozygotes susceptible (SS) (Figure 2a). On the other hand, out of the 300 unfed mosquitoes randomly selected for genotyping, 5% (15/300), 32% (62/300) and 63% (189/300), were homozygote resistant, heterozygotes and homozygote susceptible, respectively (Figure 2a). However, the distribution of L119F genotypes was not statistically different between blood-fed and unfed mosquitoes (χ^2^=0.63, *p*=0.7). Furthermore, estimation of odds ratio. (OR=1; CI 95%: 0.5-2.0; *p* = 0.6) showed overall that mosquitoes bearing the 119F-R resistant allele have the same chance to have a successful blood feeding than those with the 119F-S susceptible alleles (Table 2). This suggests that the ability to take blood is not impacted by the *L119F-GSTe2* mutation in *An. funestus*.

**Figure 2:**
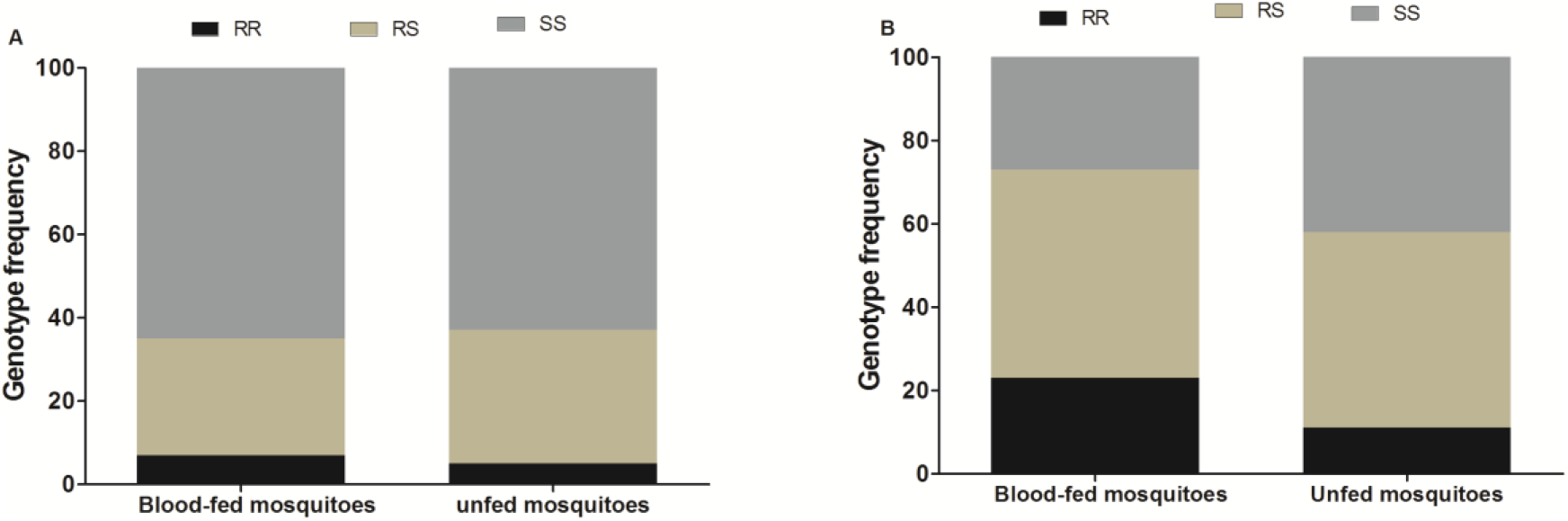
Distribution of *L119F-GSTe2* (A) and *CYP6P9a*-R (B) genotypes between blood-fed and unfed *An. funestus* mosquitoes.

**Table 2:** Assessment of the association of *L119F-GSTe2* and *CYP6P9a*-R mutations with *An. funestus* mosquito blood feeding

| Genotypes | *L119F-GSTe2* |  | *CYP6P9a-R* |  |
| --- | --- | --- | --- | --- |
|  | Odds ratio | *p*-value | Odds ratio | *p*-value |
| RR vs SS | **0.7 (0.2-2.4)** | **0.4** | **3.3 (1.4-7.7)** | **0.01** |
| RS vs SS | **1.1 (0.62-2.1)** | **0.4** | 1.7 (0.9-3.1) | **0.1** |
| RR vs RS | **0.6 (0.2-2.3)** | **0.3** | **1.9 (0.9-4.4)** | **0.1** |

SS: homozygote susceptible; RR: homozygote resistant; RS: heterozygote; \* significant difference $p < 0.05$ .

|  |  |  |  |  |
| --- | --- | --- | --- | --- |
| R vs S | 1 (0.5-2.0) | 0.6 | 1.8 (1.1-3.2) | 0.04 |

#### CYP6P9a-R

Among a total of 273 mosquitoes that were offered a blood meal 140 successfully fed (51.3%) whereas, 133 did not. Out of the 140 mosquitoes that blood-fed, 134 were successfully genotyped for *CYP6P9a*-R allele revealing that 23% (31/134), 50% (67/134) and 27% (36/134) were homozygote resistant CYP6P9a-RR, heterozygotes CYP6P9a-RS and homozygote susceptible CYP6P9a-SS, respectively (Figure 2b). Among the unfed mosquitoes, 11.3% (15/133) were homozygote resistant CYP6P9a-RR, 47.4% (63/133) heterozygotes and 41.3% (55/133) were homozygote susceptible CYP6P9a-SS. The estimation of odds ratio showed that homozygote resistant CYP6P9a-RR mosquitoes are significantly more able to blood feed that homozygote susceptible (OR = 3.33; CI 95%: 1.4 - 7.7; *p* =0.01). No difference was observed between heterozygote and homozygote resistant CYP6P9a-RR (OR= 1.9, 95%CI: 0.9-4.4; p=0.1) neither with homozygote susceptible CYP6P9a-SS (OR= 1.7, 95%CI: 0.9-3.1; p=0.1) mosquitoes (Table 2). Moreover, it was overall observed that mosquitoes with the *CYP6P9a*-R resistant allele have a greater chance to blood feed than those bearing the susceptible allele (OR = 1.9; CI 95%: 1.03-3.2; *p* =0.04) (Table 2).

### Impact of *L119F-GSTe2* mutation on probing and blood feeding duration

Out of the 120 mosquitoes that were individually filmed to assess the influence of insecticide resistance genes on the probing and feeding duration, 7 (6.14%), 40 (35.08%) and 67 (58.77%) were genotyped as homozygous resistant 119F/F-RR, heterozygotes L119F-RS and homozygotes susceptible L/L119, respectively. Overall, regardless of the genotype, the median value of mosquito’s probing duration was 49.5 seconds (minimum = 4s and maximum = 290s). No difference was observed in the probing time of resistant mosquitoes 119F/F-RR (Median = 53 seconds) and heterozygotes L119F-RS (Median = 52s) compared to the homozygote susceptible L/L119 (Median = 52s).

Regarding the blood feeding duration, it was observed that the median and mean time for a mosquito to have a full blood meal was 249.5 seconds and 303 ± 181 seconds respectively, with a minimum = 68 seconds and a maximum =772 seconds. The feeding duration was longer (median=269s) in L/L119 mosquitoes compared to L119F-RS (229.5s) and 119F/F-RR (214s) but the difference was not statistically significant (*p*=0.19, Kruskal-Wallis test).

### Impact of *L119F-GSTe2* and *CYP6P9a-R* mutations on the blood meal size of *An. funestus*

#### L119F-GSTe2

From 457 individuals that took a full blood meal it was observed that the average weighted blood meal of a mosquito regardless of the *L119F-GSTe2* genotype was 3.4 μl/mg (minimum = 1.2 μl/mg; maximum = 9.2 μl/mg). However, the weighted blood meal was not significantly different (P=0.17; Kruskal-Wallis test; Figure 3a) in homozygote susceptible L119-SS (3.0μl/mg) compared to homozygote resistant L119-RR (2.8μl/mg) and heterozygote L119F-RS (3.3μl/mg) mosquitoes. This result suggests that the *L119F-GSTe2* mutation may not affect the volume of blood meal ingested by *An. funestus*.

**Figure 3:**
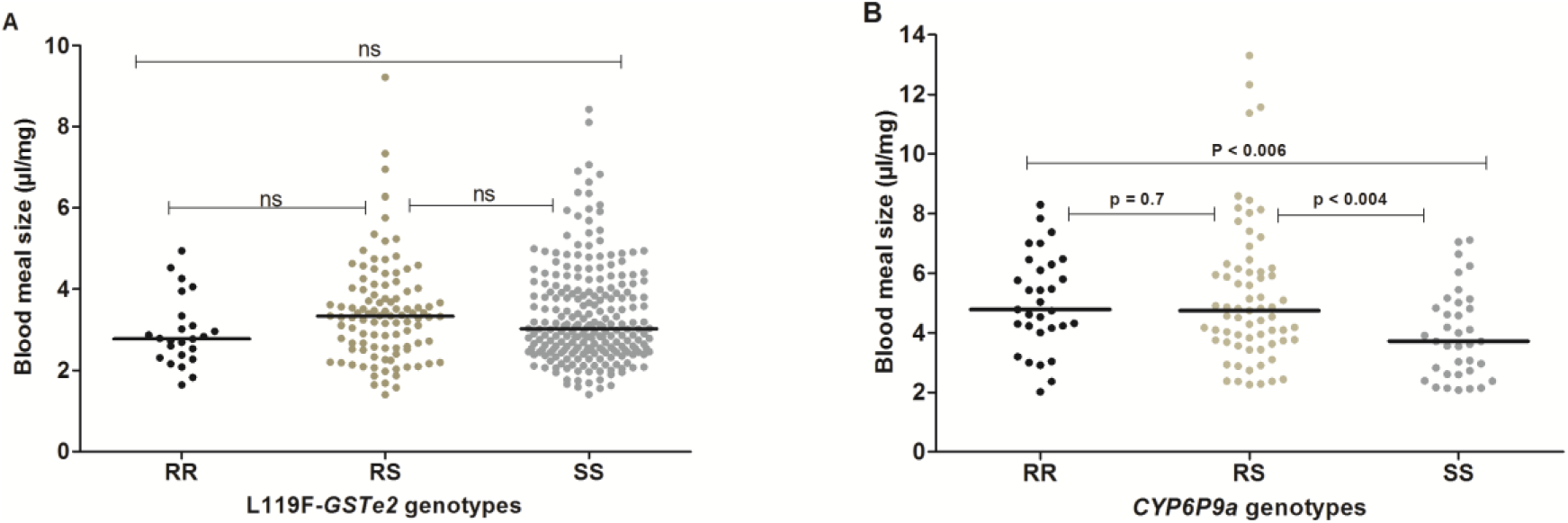
Blood meal size of *An. funestus* mosquitoes according to their *L119F-GSTe2* (A) and *CYP6P9a*-R (B) genotypes.

#### CYP6P9a-R

The influence of the *CYP6P9a*-R mutation on the volume of blood meal taken by *An. funestus*, was assessed using the 134 blood fed mosquitoes that were successfully genotyped for *CYP6P9a*-R allele. Overall, irrespective of the genotype, the mean weighted blood volume ingested by a mosquito was 4.8 ± 2 μl/mg (minimum = 2 μl/mg; maximum = 13.3μl/mg). However, the weighted blood meal volume of CYP6P9a-SS mosquitoes (Median = 3.71μl/mg) was lower than that of CYP6P9a-RS (Median = 4.73 μl/mg) and of CYP6P9a-RR (Median = 4.78 μl/mg) (Figure 3; p<0.004 for RS vs SS and p<0.006 for RR vs SS, Mann-Whitney test). No difference in the volume of the blood meal was observed between CYP6P9a-RR and CYP6P9a-RS mosquitoes (P=0.7; Mann-Whitney test). This result suggests that the over-expression of *CYP6P9a* gene is associated with an increase of the volume of the blood meal ingested by *An. funestus*.

### Expression profile of *AAPP* and *D7* family salivary genes according to *CYP6P9a-R* genotypes

Due to the association observed between the *CYP6P9a*-R genotypes and blood feeding, an attempt was made to assess whether the genotypes of this gene could impact the expression profile of key salivary genes. Analysis of the expression level of AAPP and 4 members of the D7 family salivary genes (D7r1, D7r2, D7r3 and D7r4) did not show a significant difference in expression in homozygote resistant (CYP6P9a-RR) and heterozygote (CYP6P9a-RS) mosquitoes in comparison to homozygote susceptible genotype (CYP6P9a-SS) (Figure 4, Table 2) with average fold-change for all these genes lower than 1.5. This result suggests that *CYP6P9a*-R genotypes do not influence the expression profile of both AAPP and D7 family genes in the salivary glands of *An. funestus* mosquito.

**Figure 4:**
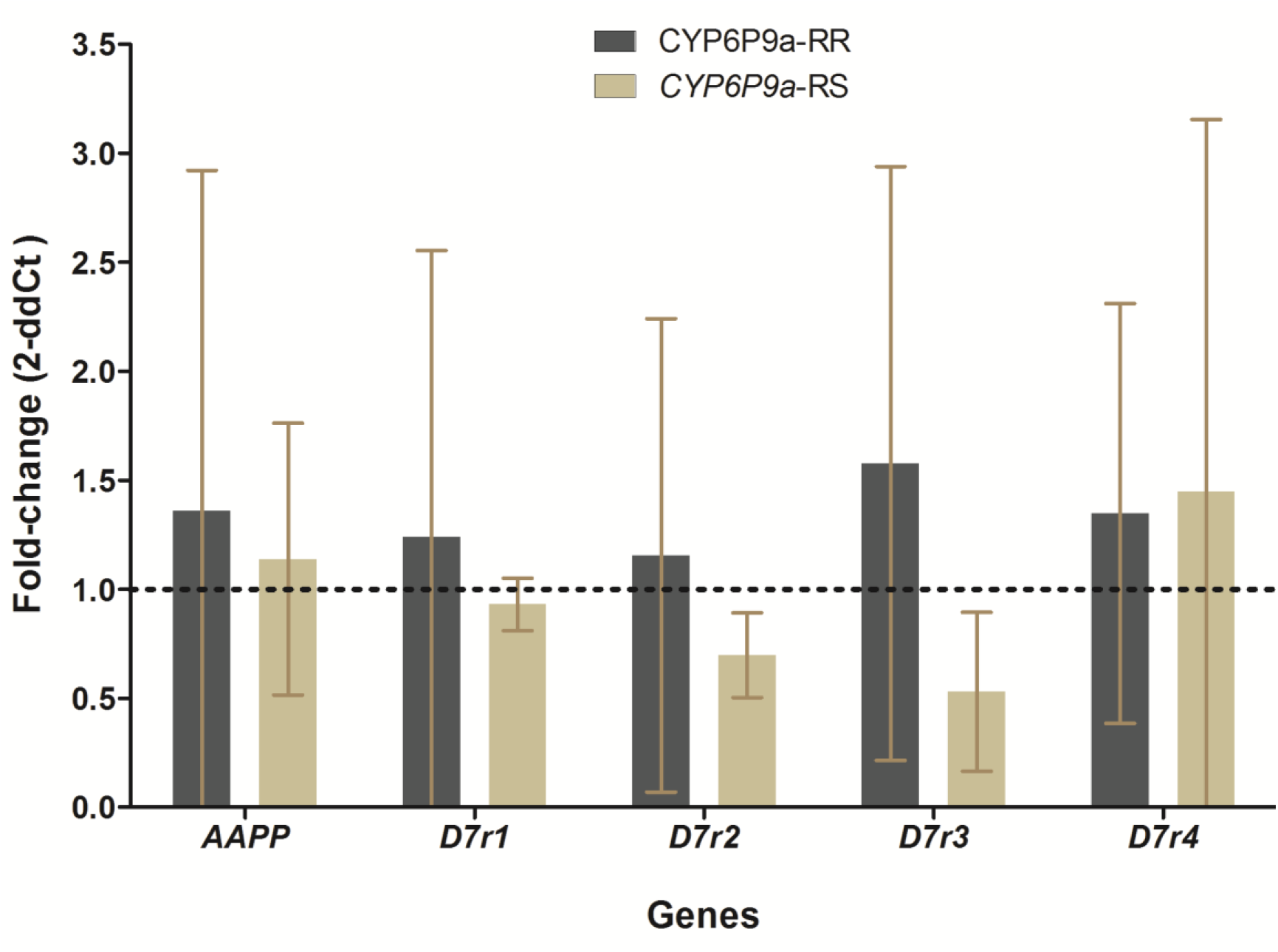
Expression level of AAPP and some members of D7 family genes in CYP6P9a-RR and CYP6P9a-RS mosquitoes in comparison with *CYP6P9a* susceptible mosquito. The dotted line indicates genes expression level in *CYP6P9a* susceptible mosquito used as standard.

## Discussion

Due to the absence of markers, the impact of metabolic resistance on life traits of *Anopheles* mosquitoes has been poorly elucidated. Recently, mutations in the *GST* epsilon 2 and in the promoter region of the cytochrome P450 *CYP6P9a*, were described as robust molecular markers for tracking metabolic resistance in pyrethroids resistant populations of *An. funestus* [22,24]. Using these two key markers, this study assess the impact of GST- and P450-based metabolic resistance to pyrethroids on the feeding process and blood meal volume of *An. funestus*.

### Impact of metabolic resistance on blood feeding success

The present study revealed that *CYP6P9a* but not the *L119F-GSTe2* mutations could impact the blood feeding success of *An. funestus* mosquito as possessing the *CYP6P9a* resistant allele increased the likelihood of being successful in blood-feeding. Such selective advantage of *CYP6P9a* resistance allele was also previously reported in a semi-field study in experimental hut trial which observed that homozygous CYP6P9a-RR mosquitoes were significantly more likely to blood feed than susceptible SS [24]. This result suggests that *CYP6P9a* -mediated metabolic resistance might influence the ability of *An. funestus* mosquito to blood feed. In contrast, the absence of association observed here for the *L119F-GSTe2* mutation needed to be confirmed by further studies as the low sample of L119F-RR homozygote resistance mosquitoes might have biased our analysis. Nevertheless, the mechanism whereby *CYP6P9a*-R resistant allele could impact mosquito feeding is unknown and was not investigated in the present study. One hypothesis to explain this association could be related to the motivation of mosquito to blood feed. In fact, it has been reported that some mosquito individuals which emerged with insufficient teneral reserves required to use their initial blood meal not to develop eggs during their first gonotrophic cycle, but to compensate for insufficient teneral reserves [33–35]. This phenomenon is mostly observed in smaller female mosquitoes which emerge with insufficient reserve [2]. Thus, we can presume that *CYP6P9a* resistant mosquitoes which were found significantly smaller than susceptible in the present study were more motivated to blood feed as they were probably the ones requiring more to compensate for their insufficient teneral reserve. Further studies investigating the impact of insecticide resistance on the motivation of *An. funestus* mosquito to blood feed would probably help confirming this hypothesis.

### Impact of metabolic resistance on probing time and feeding duration

The influence of metabolic resistance on probing time and feeding duration was assessed in the present study only for *L119F-GSTe2* mutation. Results revealed no significant impact of this metabolic resistance gene on the time spent by a mosquito to probe. The absence of impact of insecticide resistance on mosquito probing time was also reported for the *knock-down (kdr)* resistance gene in *Anopheles gambiae* with no difference in the probing time noticed between genotypes (RR, RS and SS) after exposure to untreated and insecticide-treated net (Diop et al, 2019). This seems to indicate that insecticide resistance might not impact the probing duration of *Anopheles* mosquito during blood feeding. However, this hypothesis must be taken with caution as, to our knowledge, in the exception of the present study as well as the one of Diop et *al*, very little information is available on the impact of insecticide resistance on the probing time during mosquitoes blood-feeding. In the other hand, even if the difference was not statistically significant, mosquitoes possessing an *119F-GSTe2* resistant allele (both homozygous and heterozygous) spent less time taking their blood meal than susceptible. This corroborate with observation previously made for *kdr* mutation in *An. gambiae* with lower feeding duration for homozygous resistant mosquitoes than heterozygote and homozygous susceptible [11]. The non-significant result observed may be due to the low number of resistant mosquitoes in the present study. However, it could be hypothesized that this result indicates that *L119F-GSTe2* mutation might confer an advantage to homozygous resistant mosquitoes as it was previously reported that fast feeding reduces the risk to be kill because of the host defensive behaviour [11,36].

### Impact of metabolic resistance on blood meal volume

In the present study, we observed that volume of blood ingested by a mosquito during a single blood feeding was associated with the genotypes of the P450 *CYP6P9a* but not with the *L119F-GSTe2*-based metabolic resistance. This suggests that mechanisms involved in metabolic resistance to pyrethroids in *An. funestus* might differently influence mosquito life-traits. However, as already discussed above, we cannot exclude that the absence of influence observed for *L119F-Gste2* gene might also be related to the low number of L119F-RR mosquitoes used in the present study. This latter hypothesis seems moreover reinforced by results of previous studies showing *L119F-GSTe2* mutation [23] and *CYP6P9a*-R resistance gene [37] affecting *An. funestus* fecundity in the same way. The positive association between *CYP6P9a*-R resistant allele and the volume of blood meal is a bit surprising knowing that activity of P450 monoxygenases as well as blood meal digestion, have been reported to generate an excess production of reactive oxygen species (ROS) increasing oxidative stress which could induce several damages in mosquito organism even death [38,39]. In fact, because the *CYP6P9a*-R resistant allele was recently reported to be negatively associated with the fecundity of *An. funestus* [37], we were expecting to see *CYP6P9a* resistant mosquitoes taking lower blood meal than susceptible to reduce negative effects of oxidative stress. This suggests that association between the *CYP6P9a*-R resistant allele and mosquito’s blood meal size could be an indirect consequence of some other physiological activities. For instance, because *CYP6P9a* resistant mosquitoes were significantly smaller than susceptible, and knowing that it was demonstrated that the amount of teneral reserves is proportional to the body size of mosquito [2], we can presume that the high blood meal volume ingested by CYP6P9a-RR mosquitoes might be due to a high need for these mosquitoes to compensate the limited teneral reserves after their emergence. In this case the association observed here could be an indirect consequence of the negative influence of *CYP6P9a*-R resistant allele recently observed on the larval development of *An. funestus* [37] resulting to a small body size, and by consequence to insufficient teneral reserve for resistant mosquitoes. Indeed, it was demonstrated that nutritional environment experienced by larvae strongly influences adult fitness-related traits such as body size, teneral metabolic reserves [2,30,40]. However, our finding did not corroborate with the positive association previously reported between the volume of ingested blood meal and mosquito body size [2]. Further studies will help elucidate the underlying reason of this correlation between *CYP6P9a* genotypes and blood meal size.

### Influence of CYP6P9a-R resistant allele on salivary gland genes expression

To obtain a successful blood meal, a female mosquito must balance the risk of death caused by host defensive behavior against the benefits to feed on a host species that maximize fertility [41]. Salivary components permit mosquitoes to reduce their engorgement time and increase their likelihood of survival [5]. In the present study, we assessed the level of expression of genes encoding for some salivary proteins known to be involved on blood intake process of mosquitoes such as, AAPP and members of D7 family proteins [6,42,43]. The comparative analysis of the expression level of these genes between *CYP6P9a* genotypes showed no significant difference between mosquitoes bearing the resistant allele and those with the susceptible one. This result suggests that the expression of AAPP and D7 family salivary genes is not associated to the *CYP6P9a* mutation. This observation is intriguing as some salivary genes such as D7 family genes were previously reported to be over-expressed in resistant *An. funestus* mosquito compared to susceptible strain [22,44–47]. The absence of significance of genes differential expression in the present study could be explained by the fact that, our analyses in this study were performed on mosquitoes obtained after reciprocal crosses between two different strain and therefore sharing the same background, while other studies compared insecticide resistant field mosquitoes and susceptible laboratory strains. The absence of influence of the *CYP6P9a* gene in the expression level of salivary genes involved on the blood feeding process observed in the present study appears to indicate that the association found between this gene and the size of blood meal taken by *An. funestus* mosquito might not be related with the expression of these salivary gland genes encoding proteins involved in the blood meal process.

This study revealed that *GSTe2*-mediated resistance has no effect on the blood meal intake of *An. funestus* mosquito, whereas *CYP6P9a*-based resistance to pyrethroids is associated with a feeding success and a highest blood meal size. However, this influence on *Anopheles funestus* blood meal intake is not associated with differential expression of major salivary proteins involved in blood feeding. Given the rapid growth of insecticide resistance, it would be interesting to study how this association could impact the fecundity and the vectorial capacity of *An. funestus* mosquitoes.

## Author Contributions

E.E.N and C.S.W conceived the study; EEN, L.N, C.N SK and C.S.W designed the study; E.E.N, .L.N, A.B, T.A and M.T carried out the sample collection; L.N, A.B, and T.A reared and maintained the strain in the insectary; E.E.N, L.N A.B, and A.T performed blood feeding experiments. L.N, T.A and M.T performed the molecular analyses; E.E.N, L.N, A.B, M.T and C.N analyzed the data; E.E.N, L.N and C.S.W. Emmanuel wrote the manuscript. M.T C.N and S.K reviewed the manuscript. All authors approved the manuscript.

## Ethical approval and consent to participate

Ethical clearance was obtained from the National Ethics Committee of Cameroon’s Ministry of Public Health (N°2018/04/1000/CE/CNERSH/SP) in conformity to the WMA Declaration of Helsinki. Informed verbal consent was obtained from household owners for using their houses for mosquito collection.

## Funding

This study was funded by a Wellcome Trust Training fellowship (109930/Z/15/Z) awarded to ELANGA N’DILLE Emmanuel.

## Conflicts of Interest

The authors declare no conflicts of interests.

